# Functional analysis of the epilepsy gene *Pcdh19* using a novel conditional GFP-reporter mouse model

**DOI:** 10.1101/2024.03.14.584898

**Authors:** Stefka Mincheva-Tasheva, Michaela Scherer, Louise J. Robertson, Sandra Piltz, Julien Bensalem, Daniel T. Pederick, Paul Q. Thomas

**Author notes:** Corresponding authors: Stefka Mincheva-Tasheva and Paul Q. Thomas.

## Abstract

PCDH19 is a cell adhesion molecule belonging to the delta2-protocadherin subfamily that plays a critical role in brain development, neuronal migration, synaptic organisation, and neural circuit formation. Mutations in *PCDH19* cause PCDH19-clustering epilepsy, an infantile-onset disorder characterized by seizures and intellectual disabilities. Despite the increasing development of constitutive cellular and murine models to investigate the effects of *Pcdh19* knockout on cell–cell interactions and cellular function, the spatiotemporal consequences of its loss remain poorly understood. To address this gap, we generated and validated a novel conditional *Pcdh19* knockout mouse model incorporating a GFP reporter (*Pcdh19*-cKO-GFP), enabling cell type-specific and temporally controlled gene deletion and direct visualization of recombination events. Using a neuronal *Syn1*-Cre driver, we demonstrate that targeted deletion of *Pcdh19* in neurons results in altered hippocampal neurogenesis and mouse behaviour. We further demonstrate the versatility of this model using a doxycycline-inducible Cre system, enabling temporally controlled deletion of *Pcdh19* and the modelling of disease-relevant phenotypes. Finally, we validate adeno-associated viral (AAV) vector-mediated recombination as a strategy for precise postnatal manipulation of *Pcdh19* expression. Collectively, this *Pcdh19*-cKO-GFP model provides a powerful and flexible genetic tool to interrogate the cell type specific and temporary regulated functions of PCDH19 in the developing and postnatal brain under physiological and disease conditions.

## Introduction

PCDH19 is an adhesion molecule that belongs to the delta2-protocadherin subfamily of the cadherin family, consisting of six tandemly arranged extracellular cadherin repeats (encoded by exon 1), a single transmembrane domain, and an intracellular C-terminal tail with two conserved motifs (CM1 and CM2) (Kolc et al. 2019). In vertebrates, PCDH19 plays a critical role in cellular processes such as neuronal differentiation, axon guidance, dendritic arborization, and cell signalling (Emond, Biswas, and Jontes 2009), (Bassani et al. 2018), (Borghi et al. 2021), (Mincheva-Tasheva et al. 2021). The mechanisms through which PCDH19 executes these functions are based on its role in cell-cell adhesion interactions and gene regulation (Weiner and Jontes 2013), (Pham et al. 2017), (Pederick et al. 2018), (Gerosa et al. 2019), (Gecz and Thomas 2020), (de Nys, Kumar, and Gecz 2021), (Gerosa et al. 2022), (de Nys et al. 2024), (Newbold et al. 2026), (Mincheva-Tasheva et al. 2021), (Hoshina et al. 2021). Genomic variants in the X-linked *PCDH19* gene cause PCDH19-clustering epilepsy (PCDH19-CE), an inherited epileptic syndrome characterized by variable seizures and incompletely penetrant intellectual disability (Dibbens et al. 2008). Interestingly, PCDH19-CE has a unique pattern of inheritance whereby heterozygous females are affected whereas hemizygous (transmitting) males are asymptomatic (Dibbens et al. 2008). Nearly 80% of PCDH19-CE disease variants localise to Exon 1, underscoring the importance of the cell-cell adhesion function of this domain in normal brain development and its disruption in disease pathogenesis (Kolc et al. 2019).

In the last decade, we (Pederick et al. 2016), (Pederick et al. 2018), (Homan et al. 2018), (Mincheva-Tasheva et al. 2021), (de Nys et al. 2024), (Mincheva-Tasheva et al. 2024) and others (Hayashi et al. 2017), (Galindo-Riera et al. 2021), (Borghi et al. 2021), (Niu et al. 2024), (Ziobro et al. 2025) have generated several murine (KO, conditional KO, knock in) and cell models to investigate PCDH19 biology. These models have provided valuable insights into the consequences of mosaic *Pcdh19* elimination, including altered neuronal development, impaired cell–cell interactions, disrupted synaptic function, and network abnormalities. However, constitutive knockout approaches inherently lack temporal and spatial control of gene deletion, limiting differentiation between developmental or postnatal functions and cell population-specific analysis. Furthermore, given that PCDH19 expression is dynamically regulated throughout brain development constitutive models cannot readily address how the timing and location of gene loss influence disease pathogenesis. While robust *Pcdh19* expression is observed from early embryogenesis until the first week/two years of life in mice and humans, respectively, expression gradually decreases through adulthood in both species (Gaitan and Bouchard 2006), (Herring et al. 2022). These changes in the expression pattern suggest that it may have a tightly regulated role in brain development and physiology (Pederick et al. 2016), (Pederick et al. 2018), (Hoshina et al. 2021), (Galindo-Riera et al. 2021). These observations highlight the need for a conditional genetic model that enables precise control of *Pcdh19* deletion in defined cell populations and at selected developmental stages with precise visualisation of the recombinant cells.

Here, we describe the generation and validation of a novel conditional *Pcdh19* knockout mouse carrying an integrated GFP reporter (*Pcdh19*-cKO-GFP). This model enables cell type-specific and temporally controlled deletion of *Pcdh19* and direct visualization and characterization of Cre-recombined cells *in vivo*. Through neuronal elimination of *Pcdh19*, using the *Syn1*-Cre mouse line, we demonstrate the impact of *Pcdh19* deletion on adult neurogenesis, and behaviour. We further establish its versatility by employing a doxycycline-inducible Cre system to achieve stage-specific deletion during embryonic development, enabling temporal modelling of PCDH19-associated pathology. Finally, we demonstrate the feasibility of adeno-associated virus (AAV)-mediated Cre recombination for targeted postnatal deletion of *Pcdh19*. Altogether our observations demonstrate the feasibility of using the *Pcdh19*-cKO-GFP to investigate the impact of cell and stage-specific *Pcdh19* elimination on mouse brain physiology and pathology.

## Results

### Generation and validation of a novel *Pcdh19-*cKO-GFP mouse model

Conditional knockout mouse models are well suited for investigating cell type–specific and spatiotemporal *in vivo* functions of disease-causing genes. To explore their utility in the context of epilepsy-causing gene *PCDH19*, we generated a conditional floxed *Pcdh19* mouse model (*Pcdh19*-cKO-GFP) where Exon 1 of the gene is flanked by LoxP sites in the 5’UTR and Intron 1. In addition, a GFP reporter cassette was inserted immediately downstream of the 3’ LoxP site followed by Poly(A) signal (Fig. 1A). Upon Cre-mediated recombination, excision of Exon 1 generates a *Pcdh19* knockout (KO) allele and brings GFP expression under the control of the endogenous promoter to mark *Pcdh19*-null (KO) cells. To validate the genetic modification, diagnostic PCRs were performed to confirm the presence of the knock-in allele (Fig. 1B) and the result was further corroborated by droplet digital PCR (ddPCR) (Supp. Fig. 1B).

**Figure 1.**
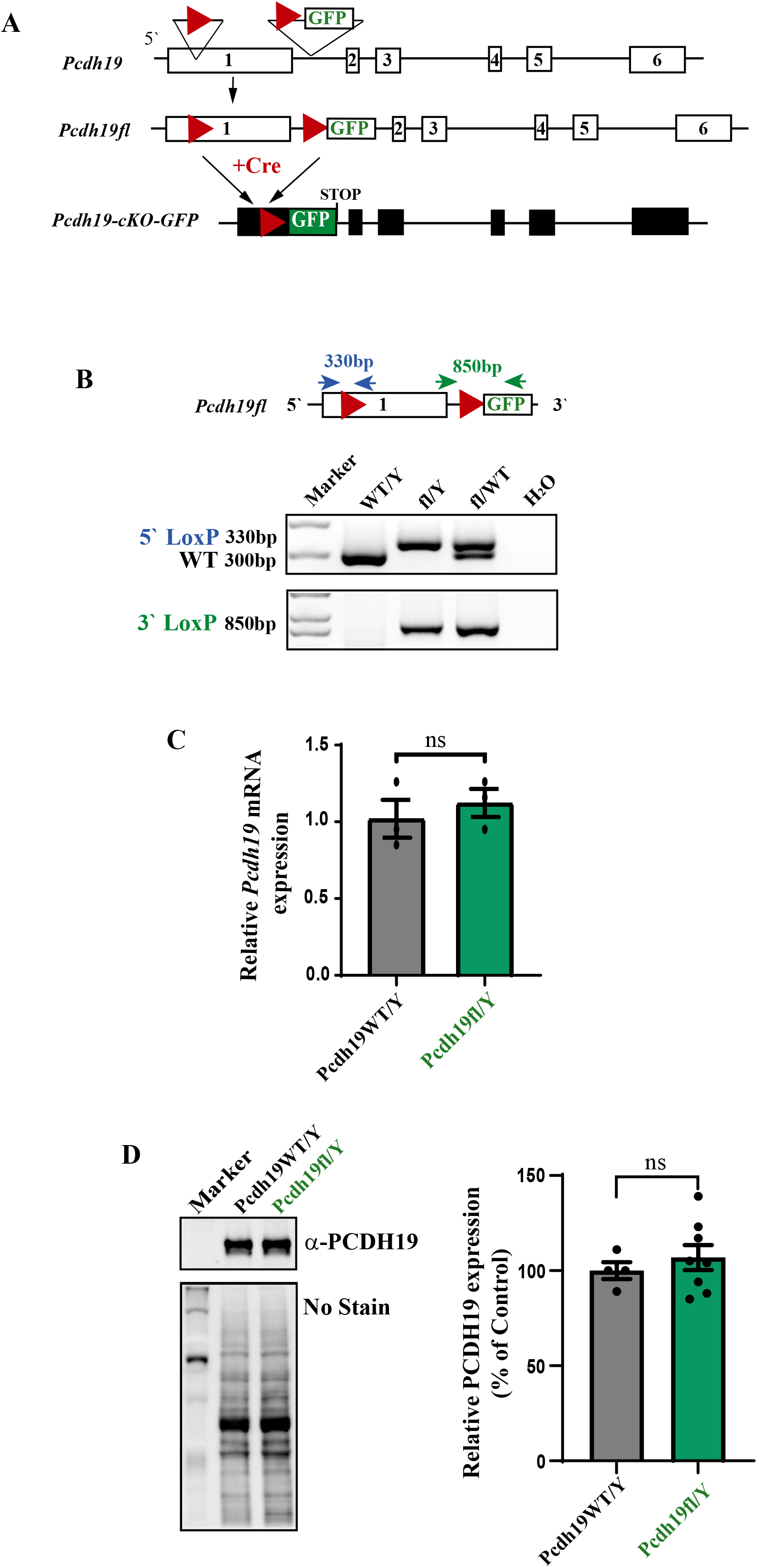
Generation of *Pcdh19*-cKO-GFP mice. (**A**) Schematic of *knock-in mouse* generation. Red arrowheads indicate the 3’ and 5’ LoxP sites. Cre-recombination leads to Exon1 deletion, GFP reporter expression and the stop codon after GFP cassette prevents the expression of the remaining exons (2-6). **(B)** PCR was used to confirm the presence of the 5′ LoxP site (left) and 3′ Lox P site and GFP cassette (right). Primers sites are indicated with blue arrowheads. (**C**) RT-qPCR of *Pcdh19* relative expression in *Pcdh19*^WT/Y^ and *Pcdh19*^*fl/Y*^ mice. mRNA was normalised to *Gapdh* mRNA expression level (n=3). **(D)**. Western Blotting of *Pcdh19*^*WT/Y*^ (WT/Y) and *Pcdh19*^*fl*/Y^ mouse brains at P2 using anti-PCDH19 and No-stain loading control. The graph represents the relative quantification of PCDH19 level in P2 *Pcdh19*^*fl/Y*^ and *Pcdh19*^*WT/Y*^ mouse brains normalised to the No stain loading control (n=4 and n=8, respectively). All data is presented as mean ± SEM.

Next, we investigated endogenous *Pcdh19* mRNA and protein levels in the cKO-GFP mouse model (in the absence of Cre recombinase). RT-qPCR and western blot analysis showed no significant difference between *Pcdh19*^*fl/Y*^ and *Pcdh19*^*WT/Y*^ brain samples confirming that the genetic architecture of the *Pcdh19*-cKO-GFP mouse model does not affect endogenous *Pcdh19* expression (Fig. 1C&D).

### Effect of neuron-specific elimination of *Pcdh19* on brain physiology

Several studies using *Pcdh19* knock-in and knock-out mouse models have shown extensive expression of *Pcdh19* in cortical and hippocampal neurons during development and postnatal stages (Pederick et al. 2018), (Rakotomamonjy et al. 2020), (Mincheva-Tasheva et al. 2021), (Hoshina et al. 2021), (Galindo-Riera et al. 2021), (Mincheva-Tasheva et al. 2024), (Lim et al. 2019). However, how cell-specific *Pcdh19* deletion impacts brain physiology is poorly understood. We investigated the efficiency and consequences of eliminating *Pcdh19* in neurons using a mouse line in which Cre is expressed in neurons from the rat Synapsin1 (*Syn1*-Cre) promotor (Zhu et al. 2001). To avoid ectopic expression of Cre in the male germline we employed a breeding strategy where the Cre-recombinase allele is transmitted only from the female dams (Rempe et al. 2006). To verify that the cells undergoing Cre recombination were of neuronal origin, we performed immunostaining using an antibody against the neuronal nuclear protein (NeuN), a marker expressed by nearly all post-mitotic neurons in cortex and hippocampus (Mullen, Buck, and Smith 1992). Immunofluorescent analysis of *Pcdh19*^*fl*/Y^; *Syn1*-Cre mice revealed prominent GFP expression in mature (NeuN positive) neurons of the brain, validating the activity of the null cell reporter cassette (Fig. 2A&B). The distribution pattern of PCDH19 KO cells in cortex and hippocampus of *Pcdh19*^*fl*/Y^;*Syn1*-Cre mice was similar to that observed in previously reported constitutive *Pcdh19-*KO and *Pcdh19*-Tag mouse models (Pederick et al., 2016), (Hayashi et al. 2017) (Pederick et al. 2018) confirming that the majority of the cells expressing *Pcdh19* are of neuronal origin. Next, we determined the PCDH19 level in cortex and hippocampus (Mullen, Buck, and Smith 1992). Immunofluorescent analysis of *Pcdh19*^*fl*/Y^; *Syn1*-Cre mice after Cre–mediated recombination. Western blot analysis revealed a 40% reduction in PCDH19 protein levels compared to controls (Fig. 2C), confirming Cre-mediated excision of the *Pcdh19* floxed allele.

**Figure 2.**
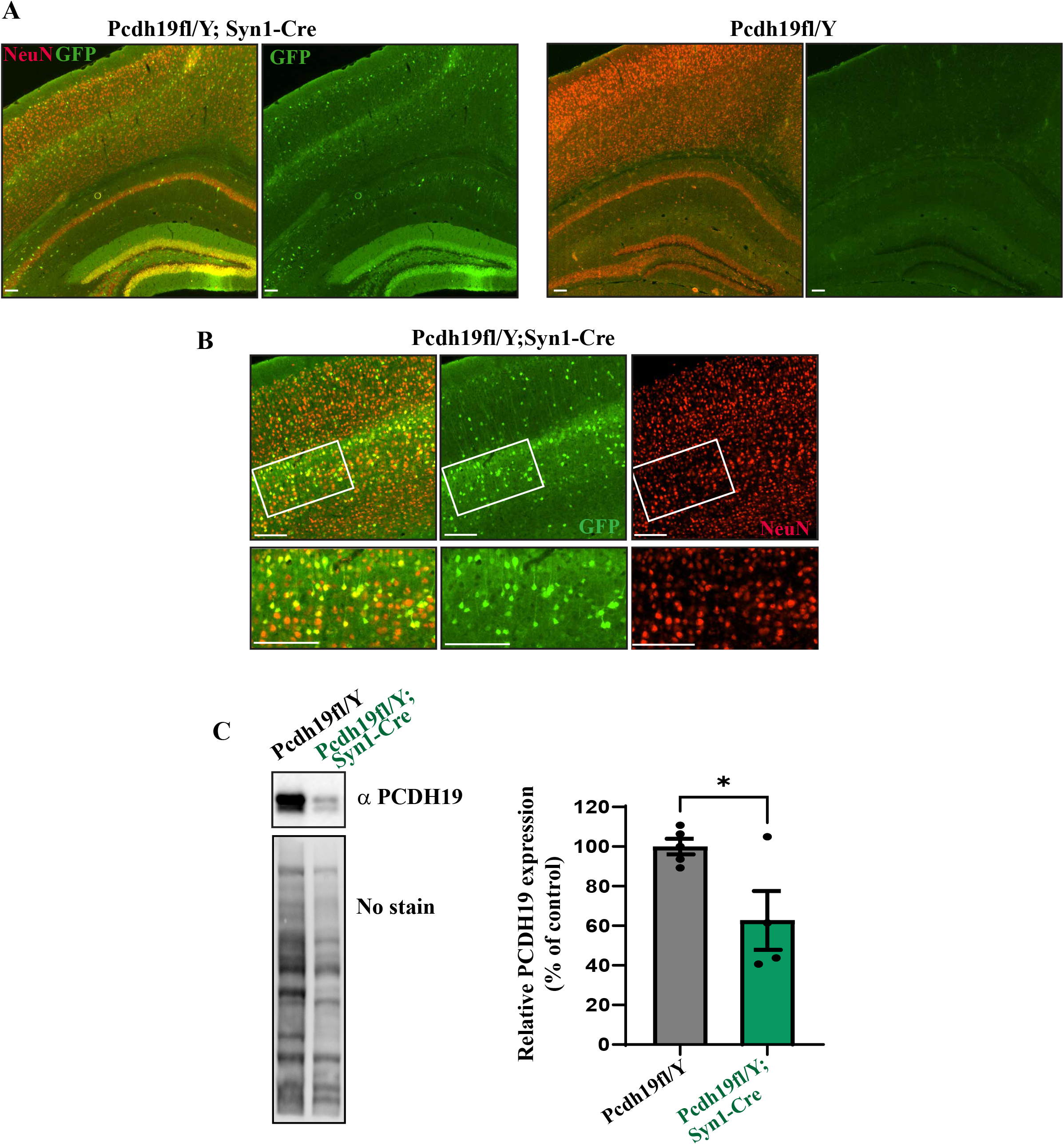
*Pcdh19* elimination in *Pcdh19*^*fl*/Y^;*Syn1*-Cre mouse model. **(A)** Immunofluorescence images of *Pcdh19*^*fl*/Y^;Syn1-Cre and *Pcdh19*^*fl*/Y^ P21 coronal brain slices stained with anti-NeuN2 (red) and anti-GFP (green) antibodies. Scale bar 100µm **(B)** Higher magnification images of cortical areas of the *Pcdh19*^*fl*/Y^;*Syn1*-Cre brains immunostained for NeuN and GFP. Scale bar 50µm. (**C**) Western blot analysis of *Pcdh19*^*fl*/Y^ and *Pcdh19*^*fl*/Y^;*Syn1*-Cre and mouse brains at P22-23 using anti-PCDH19 antibody and No stain loading control. The graph represents the quantification analysis of n≥4. All data is presented as mean ± SEM (\**p* < 0.05).

Previous studies using *in vitro* models have shown that *Pcdh19* elimination alters iPSC neurogenesis (Homan et al. 2018), (Tidball et al. 2023), (Borghi et al. 2021), (Niu et al. 2024). To investigate hippocampal neurogenesis, we assessed the expression of markers of proliferating cells (Antigen Kiel 67, Ki67) and immature neurons (doublecortin, DCX) in *Pcdh19*^*fl*/Y^;*Syn1*-Cre mice (Fig. 3A&B). We observed a significant increase in neurogenesis-associated markers following neuronal *Pcdh19* deletion compared with control (*Pcdh19*^*fl*/Y^ and *Syn1*-Cre male littermates) (Fig.3 A&B). Next, we assessed the impact of *Pcdh19* neuronal elimination on locomotor activity and exploration behaviour using the open field test (Fig. 3C). Experimental mice (*Pcdh19*^*fl*/Y^;*Syn1*-Cre) showed a significantly increased number of entries into the central arena compared with control male littermates (*Pcdh19*^*fl*/Y^ and *Syn1*-Cre) reflecting increased exploratory behaviour (Fig. 3C). Although time spent in the central arena was also elevated in the experimental group, this difference did not reach statistical significance. In contrast, total distance travelled was comparable between control (*Pcdh19*^*fl*/Y^ and *Syn1*-Cre) and experimental (*Pcdh19*^*fl*/Y^;*Syn1*-Cre/Y) mice, indicating that overall locomotor activity was unaffected. Collectively, these results indicate that neuronal *Pcdh19* loss alters neurogenesis and mouse exploration behaviour.

**Figure 3.**
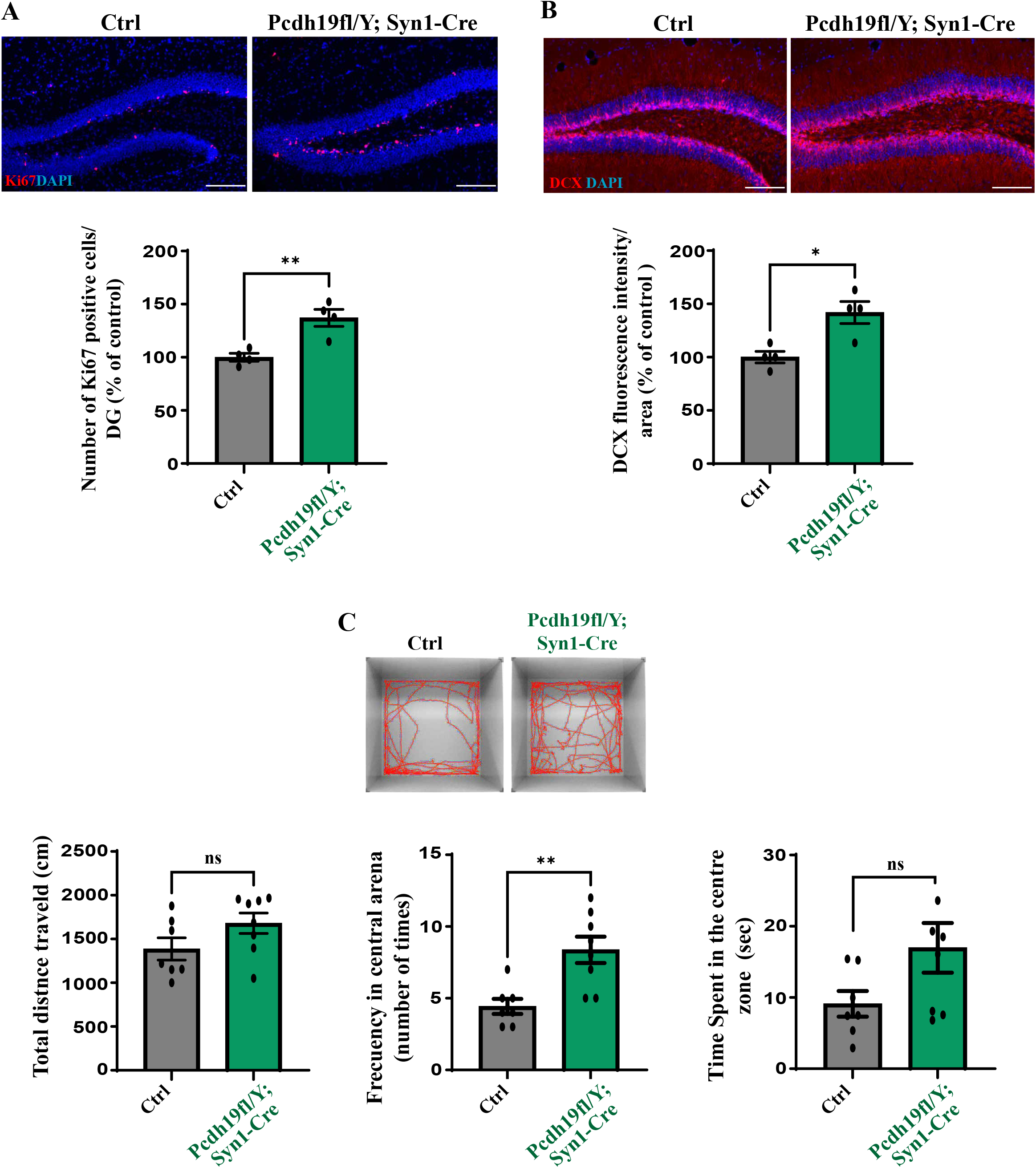
*Pcdh19* elimination in *Syn1* neurons affects neurogenesis and behaviour. **A)** Immunofluorescence images of *Pcdh19*^*fl*/Y^;*Syn1*-Cre and control (*Pcdh19*^*fl*/Y^ and male *Syn1*-Cre) brain samples stained against Ki67 (red) and DAPI (blue). The graph represents the quantitative analysis of the number of Ki67 positive cells in DG area DG of *Pcdh19*^*fl*/Y^;*Syn1*-Cre and controls (*Pcdh19*^*fl*/Y^ and *Syn1*-Cre) (n=4). **(B)** Immunofluorescence images of *Pcdh19*^*fl*/Y^;*Syn1*-Cre and male control littermates (combined *Pcdh19*^*fl*/Y^ and *Syn1*-Cre) stained with DCX (n=4). The graph represents the fluorescence intensity/area in the hippocampal DG of *Pcdh19*^*fl*/Y^;*Syn1*-Cre and controls (*Pcdh19*^*fl*/Y^ and *Syn1*-Cre). Scale bar in (A) and (B) is 100 µm. (**C**) Open field test demonstrates changes in the frequency of the central arena entries of *Pcdh19*^*fl*/Y^;*Syn1*-Cre compared to control male littermates (combined *Pcdh19*^*fl*/Y^ and *Syn1*-Cre) (n≥7). No significant changes in the total distance travel and the time spent in the central arena were observed. All data is presented as mean ± SEM (\**p* < 0.05, \*\**p* < 0.01).

### Spatiotemporal control of *Pcdh19* knockout via Dox-Cre and AAV-Cre systems

A major challenge in elucidating PCDH19-CE pathogenesis is identifying the critical developmental window during which mosaic *Pcdh19* loss alters brain development and drives neurological dysfunction. To evaluate whether the *Pcdh19*-cKO-GFP model could be used for stage-specific gene deletion, we first employed a doxycycline (Dox)-inducible Cre recombinase system (rtTA; TetO-Cre) (Perl et al. 2002). *Pcdh19*^*fl*/Y^; rtTA; Cre embryos were exposed to Dox from 3.5 dpc until 18.5 dpc and Cre-mediated recombination was assessed by diagnostic PCR (Supp.Fig. 2). Recombination, indicated by loss of the 5′ LoxP site and generation of the recombined allele, was detected exclusively in Dox-treated embryos *(Pcdh19*^*fl*/Y^;rtTA; Cre and *Pcdh19*^*fl/fl*^;rtTA;Cre), confirming inducible *Pcdh19* deletion.

To validate temporally controlled *Pcdh19* deletion, we examined GFP expression following doxycycline-induced Cre activation. In doxycycline-treated *Pcdh19*^*fl/Y*^;rtTA;Cre mice, GFP-positive (PCDH19-deficient) cells were widely distributed throughout the cortex and hippocampus (Fig. 4A). The distribution of recombined cells was consistent with that previously reported in constitutive *Pcdh19*-KO mouse models (Pederick et al. 2016), (Pederick et al. 2018), (Hayashi et al. 2017), (Galindo-Riera et al. 2021), demonstrating efficient recombination *in vivo*.

**Figure 4.**
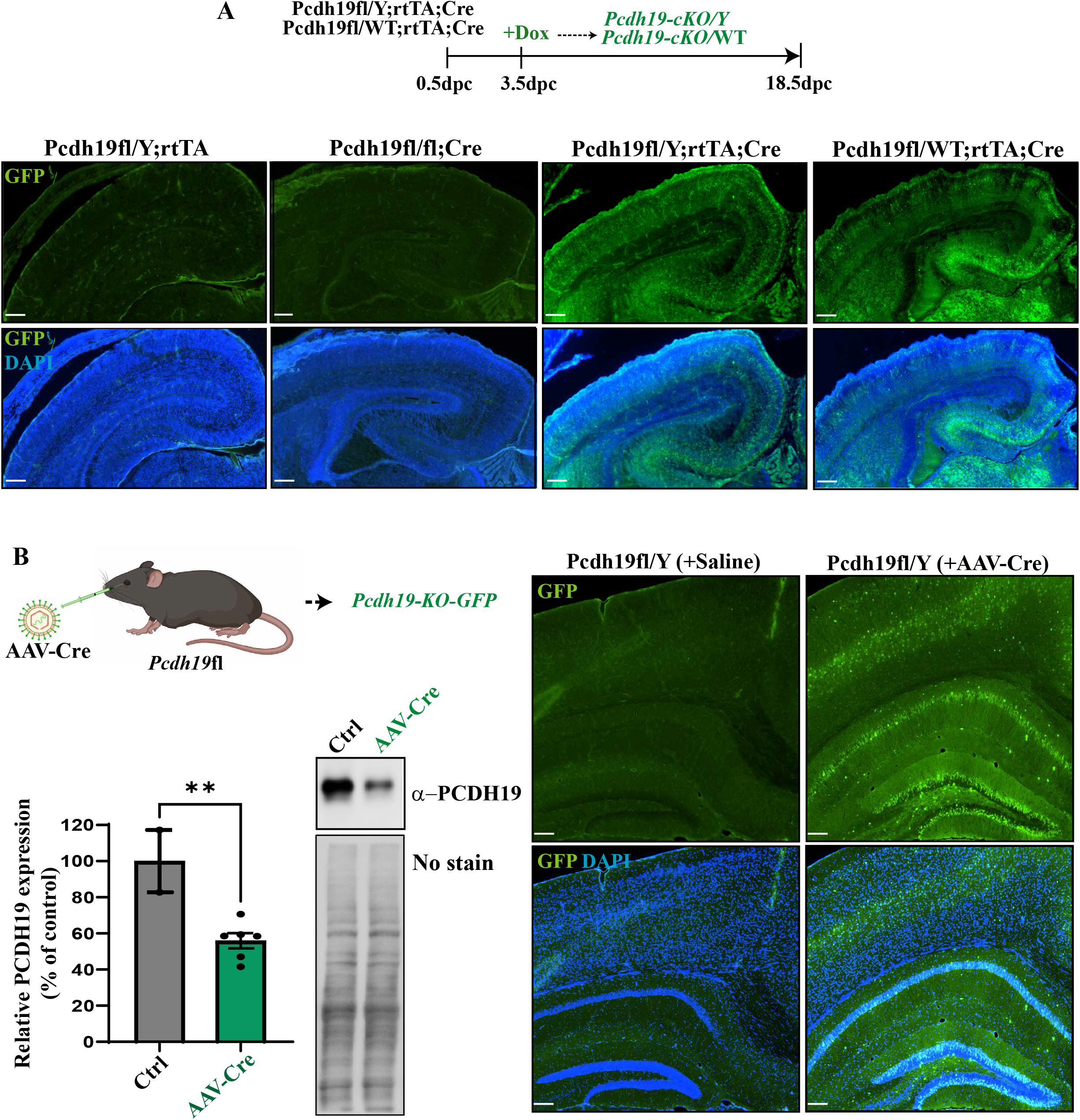
Stage-specific deletion of *Pcdh19* during embryogenesis and at postnatal stages. **A)** Schematic representation of Dox-inducible Cre activation and *Pcdh19* deletion in *Pcdh19*^*fl/*Y^;rtTA;Cre, *Pcdh19*^*fl/*WT^;rtTA;Cre mice during embryogenesis. Immunofluorescence images of coronal brain slices at 18.5dpc from *Pcdh19*^*fl/*Y^;rtTA;Cre, *Pcdh19*^*fl/*WT^;rtTA;Cre, and *Pcdh19*^*fl/fl*^;Cre, and *Pcdh19*^*fl/Y*^;rtTA control male and female mice treated with Dox from 3.5dpc to18.5dpc. Sections were stained with anti-GFP (green) antibody to indicate recombination (PCDH19-KO cells), and DAPI (blue) for nuclear staining (n≥3,). (**B)** Illustration (created with BioRender.com) of systemic retro-orbital injection (ROI) of AAV-PHP.eB-EF1a-Cre into *Pcdh19*^*fl/Y*^ mice at P15. WB analysis of control (saline) and AAV-Cre *Pcdh19*^*fl/Y*^ mice 7 days after ROI demonstrated significant reduction of PCDH19 level in the AAV-Cre transduced brains (n=6) compared to control mice (n=2). All data is presented as mean ± SEM (*t*-test, \*\**p* < 0.01). Immunofluorescence images in (B) of brain sections collected 7 days post-injection show GFP expression (green) in cortex and hippocampus (left) compared to saline-injected controls (right). Nuclei are counterstained with DAPI (blue). Scale bar in (A) and (B) is 100 µm.

We next investigated whether embryonic induction of Cre activity could reproduce the mosaic pattern of PCDH19 loss characteristic of PCDH19-CE. Dox administration at 3.5 dpc in female *Pcdh19*^*fl/WT*^;rtTA;Cre mice generated mosaic population of PCDH19-deficient (GFP-positive) cells and robust cell segregation throughout the cortex and hippocampus (Fig. 4A). Importantly, no recombination was detected in the absence of Dox (Supp.Fig. 2), confirming the strict inducibility of the system. Together, these findings validate the *Pcdh19*-cKO-GFP model as a robust platform for temporally controlled embryonic gene deletion and for investigating the developmental role of PCDH19 in brain organization.

To extend this approach to postnatal gene disruption, we next utilised systemic AAV-mediated gene delivery, a well-established method for efficient, non-invasive brain transduction in mice (Mathiesen et al. 2020). *Pcdh19*-flox mice at postnatal day 15 (P15) received a retro-orbital injection of AAV-PHP.eB-EF1α-Cre. Cre-mediated recombination was assessed seven days post-injection using immunofluorescence and western blot analysis (Fig. 4B). Robust GFP expression was observed in neurons across the cortex and hippocampus of AAV-transduced *Pcdh19*^*fl/Y*^ brains, whereas no GFP signal was detected in saline-injected controls. Consistent with efficient recombination, western blot analysis demonstrated an approximately 50% reduction in PCDH19 protein levels in AAV-Cre–treated brains compared with controls (Fig. 4B). Collectively, these findings demonstrate that the *Pcdh19*-cKO-GFP model supports efficient inducible gene deletion during both embryonic and postnatal development.

## Discussion

Over the past decade, constitutive *Pcdh19* KO mouse models have been instrumental in characterising the role of PCDH19 in brain development and disease (Pederick et al. 2016), (Hayashi et al. 2017), (Pederick et al. 2018), (Lim et al. 2019), (Galindo-Riera et al. 2021), (Hoshina et al. 2021). Although these models have facilitated mapping of PCDH19-deficient cells in the brain, they do not allow temporal or cell-type specific control over *Pcdh19* deletion. This limitation has hindered efforts to define the contribution of individual cell populations to the diverse neurodevelopmental and behavioural phenotypes associated with mosaic *Pcdh19* deletion. To date, only one conditional *Pcdh19* KO model has been reported (Giansante et al. 2023). While this represents an important advance for the field, the model does not incorporate a reporter gene to enable tracing of the identity and distribution of *Pcdh19*-KO cells. Given that not all cells in the cortex or hippocampus express *Pcdh19* (Pederick et al. 2016), (Hayashi et al. 2017), (Galindo-Riera et al. 2021), tracing the identity and distribution of recombined cells can be challenging. To extend the capabilities of existing models and provide additional resolution in mosaic contexts, we developed and validated a novel *Pcdh19*-cKO-GFP mouse model, in which GFP acts as an endogenous reporter for Cre-mediated deletion. This design enables direct visualization and tracking of *Pcdh19*-null cells, facilitating the characterization of their distribution and contribution to circuit-level and behavioural phenotypes in a mosaic context, a hallmark of PCDH19-CE.

Although *Pcdh19* is expressed in neuronal and non-neuronal brain cells (Galindo-Riera et al. 2021) in the cortex and hippocampus, the impact of *Pcdh19* deletion on distinct cell populations is yet to be determined. Here, we show a similar distribution pattern of *Pcdh19*-KO cells in the neuron-specific (*Pcdh19*^*fl*/Y^;*Syn1*-Cre) and the previously reported constitutive *Pcdh19*-KO mouse models (Pederick et al. 2016), (Hayashi et al. 2017; Galindo-Riera et al. 2021), indicating that the vast majority of the cells expressing *Pcdh19* have a neuronal origin.

While adult neurogenesis has a profound impact on learning and memory formation, increasing evidence links dysregulated adult neurogenesis to neurological disease, including epilepsy (Chen et al. 2021). Although substantial progress has been made in investigating the role of *Pcdh19* in neurogenesis using *Pcdh19-*KO mouse models and induced pluripotent stem cells (iPSCs) (Pederick et al. 2016), (Homan et al. 2018), (Borghi et al. 2021), (Tidball et al. 2023), there remains a significant knowledge gap regarding the specific cell types and brain regions that influence this process. Here we show that deletion of *Pcdh19* in *Syn1*-expressing neurons resulted in alterations in hippocampal neurogenesis during young adulthood. Furthermore, our study revealed significant changes in the frequency of entry in the central arena of the open field in *Pcdh19*^*fl*/Y^;*Syn1-*Cre mice which has been linked to a risk-taking and rapid exploration behaviour (Powell et al. 2004), (Langford-Smith et al. 2011). Together, these findings raise the possibility that altered neurogenesis contributes to the exploratory phenotype observed in *Pcdh19*^*fl/Y*^;Syn1-Cre mice, although a direct mechanistic link between these two processes remains to be established.

The dynamic and robust expression pattern of *Pcdh19* during embryonic development, followed by more restricted expression in the postnatal brain (Hayashi et al. 2017), (Pederick et al. 2018) suggests that distinct developmental windows may differ in their sensitivity to gene loss. Mutations in *PCDH19* cause PCDH19-CE in heterozygous females and mosaic males. A widely accepted pathogenic mechanism is the phenomenon of cellular interference, whereby intermixed populations of PCDH19-expressing and PCDH19-deficient cells segregate during brain development, disrupting cell-cell communication and altering neuronal network function (Depienne and LeGuern 2012), (Pederick et al. 2018), (Mincheva-Tasheva et al. 2021). Despite strong genetic and experimental support for this model, the developmental period during which mosaic *Pcdh19* deletion leads to cell segregation and CNS dysfunction remains poorly defined. The inducible Dox-dependent Cre system provides precise temporal control of recombination, allowing systematic interrogation of how the developmental timing of *Pcdh19* loss influences mosaic architecture, cellular segregation, and downstream brain phenotypes across development. Here, we show that Dox administration from 3.5 dpc efficiently induces recombination, resulting in a widespread distribution of *Pcdh19*-null cells in *Pcdh19*^*fl/Y*^ brains and the generation of mosaic *Pcdh19*-deficient cell populations with pronounced cellular segregation in heterozygous *Pcdh19*^*fl/WT*^ brains. These phenotypes closely recapitulate key features observed in constitutive *Pcdh19* models while further providing the temporal flexibility required to dissect the developmental consequences of mosaicism. Importantly, this system offers a versatile platform for inducing *Pcdh19* loss at defined developmental stages and determining further how the temporal mosaic formation influences brain organisation and function.

Viral delivery of Cre enables temporal control of gene deletion postnatally which can be adapted for widespread or region-specific recombination depending on the route of administration and vector design. Consistent with these advantages, systemic administration of AAV-PHP.eB-Cre efficiently induced recombination throughout the postnatal brain, resulting in robust *Pcdh19* depletion across cortical and hippocampal regions. These findings establish the *Pcdh19*-cKO-GFP mouse as a platform that is readily adaptable to viral-based experimental paradigms.

In conclusion, the *Pcdh19*-cKO-GFP mouse model represents a versatile and powerful tool for studying PCDH19 function in brain development, neurogenesis, and behaviour. By enabling both conditional gene deletion and direct visualisation of recombined (*Pcdh19* KO) cells, this model addresses a key limitation in the field and allows precise dissection of mosaic gene function *in vivo*. Ultimately, this versatility provides a valuable platform for defining the developmental timing and cellular consequences of *Pcdh19* loss in the brain and for dissecting the mechanisms underlying PCDH19-CE pathogenesis.

## Materials and Methods

### Experimental mice housing

All animals used for this study were housed at South Australian Health and Medical Research Institute (SAHMRI). Experiments were performed using the mouse lines listed above under the SAHMRI Animal Ethics Committee Approvals SAM20-015 and SAM24-089. The animal facility is regulated for temperature and maintained on a 12 h light–dark cycle with 24 h food and water *ad libitum*. Mice of both sexes were used for all experiments.

### Generation of conditional *Pcdh19* floxed mouse model (*Pcdh19*-cKO-GFP)

CRISPR gRNAs were generated to target the *Pcdh19* Exon1 5′UTR region with gRNA1 5’-TCACTTCCCTAGCCCCTCGC and gRNA2 targeting Intron 1: 5’-GTGAACTATAACCCTACAGC. Microinjection and gRNA production was carried out as previously described (Robertson et al. 2018). We used a two-stage process to generate the *Pcdh19*-cKO-GFP mouse model. Firstly, a single-stranded donor to introduce 3’LoxP site and the GFP cassette (eGFP followed by Poly(A) signal, flanked with FRT sites), with 80 bp upstream and 36 bp downstream homology arms was generated using a plasmid generated at Vector Builder as a template (Supp.Fig. 1A). 10 ng/μL of ssDNA donor, 25 ng/μL gRNA2 and 50 ng/uL SpCas9 mRNA was microinjected into the pronucleus of C57BL/6J zygotes, transferred to pseudo-pregnant recipients, and allowed to develop to term. Founder pups were screened by PCR amplifying the 3’LoxP site and part of the GFP region: Fwd 5’-GTCATGGTTGGGAGGGCTTG and Rev 5’-CCATGTGATCGCGCTTCTCG (knock-in band – 850 bp, whereas no band was present in WT mice). The second stage used a single-stranded DNA donor purchased from IDT, including the 5’ LoxP sequence flanked by 60bp homology arms (Supp. Fig. 1A). 100 ng/μL of ssDNA donor, 50 ng/μL gRNA1 and 100 ng/μL SpCas9 mRNA was microinjected into the pronucleus of zygotes positive for the 3’ LoxP site, transferred to pseudo-pregnant recipients, and allowed to develop to term. The presence of the 5’ LoxP site was confirmed by PCR using the following primers: Fwd 5’-CGCTGATTGGCTAGTGCCTG and Rev 5’-GGTCTCAGATGAATCGGCTC (a wild-type band of ~300 bp and a 5’ LoxP of ~330 bp) (Fig. 1B). Sanger sequencing was used to confirm the correct sequence of the knock-in allele (data available upon request). A ddPCR was used confirm the copy number of the GFP cassette in the knock-in mice (Supp. Fig. 1B).

### Generation of *Pcdh19*^*fl/Y*^*;Syn1-*Cre^*Tg/0*^

To generate *Pcdh19*^*fl/Y*^*;Syn1-*Cre^*Tg/0*^ mice, we first crossed *Syn1*-Cre females (B6.Cg-Tg (*Syn1-* Cre) 671Jxm/J) from Jax laboratory (JAX stock #003966) with *Pcdh19*^*fl/Y*^ males. The resulting offspring *Pcdh19*^*fl/*WT^;*Syn1*-Cre^Tg/0^ were bred with *Pcdh19*^*fl/Y*^ males to establish the experimental cohorts (*Pcdh19*^*fl/*Y^;*Syn1*-Cre^Tg/0^, hereafter referred to as *Pcdh19*^*fl/Y*^*;Syn1*-Cre) and male controls (*Pcdh19*^*fl/Y*^ and *Syn1*-Cre). To confirm the presence of the *Syn1*-Cre allele in these mice, we used the following primers: Transgene Fwd: 5’-CTCAGCGCTGCCTCAGTCT and Rev 5’-GCATCGACCGGTAATGCA and Internal positive control (Fwd 5’-CTAGGCCACAGAATTGAAAGATCT and Rev: 5’-GTAGGTGGAAATTCTAGCATC). Transgene = 300 bp and Internal positive control = 324 bp. The existence of the *Pcdh19* flox allele was verified as described above.

### Generation of *Pcdh19*^*fl/*Y^;rtTA^Tg/0^;TetO-7Cre^Tg/0^ and *Pcdh19*^*fl/*WT^;rtTA^Tg/0^;TetO-7Cre^Tg/0^ mouse models

To generate the male and female inducible *Pcdh19*^fl^;rtTA^Tg/0^;TetO-7Cre^Tg/0^ mouse line (referred to herein as *Pcdh19*^*fl/*Y^;rtTA;Cre and *Pcdh19*^*fl/*WT^;rtTA;Cre) we crossed the TetO-7Cre female mouse (B6.Cg-Tg(tetO-7cre)1Jaw/J) from JAX stock #006234 containing the *Pcdh19* floxed alleles with a mouse containing the rtTA transgene (rtTA^Tg/0)^.

To genotype the TetO-7CRE transgene, we used the following primers: Transgene (Fwd 5’-GCGGTCTGGCAGTAAAAACTATC Rev: 5’-GTGAAACAGCATTGCTGTCACTT) and Internal positive control (Fwd: 5’-CTAGGCCACAGAATTGAAAGATCT and Rev: 5’-GTAGGTGGAAATTCTAGCATCATCC). Transgene = 100 bp and internal positive control = 324 bp.

rtTA mice were obtained from the Australian Phenomics Facility in Canberra, Australia (C57BL/6-Col1a1tm2(tetO-RNAi:Rps19)Karl Gt(ROSA)26Sortm1(rtTA*M2)Jae/AnuApb). We used a breeding strategy to segregate the *Rsp19* shRNA transgene away from the rtTA-M2 Rosa26 transgene. To genotype the mice containing rtTA transgene only, we used the following primers: Mutant Rev: GCGAAGAGTTTGTCCTCAACC, WT Rev: 5’-GGAGCGGGAGAAATGGATATG and Fwd: 5’-AAAGTCGCTCTGAGTTGTTAT. Mutant allele PCR products were ~340 bp and WT products were ~650 bp.

### Doxycycline (Dox) treatment of *Pcdh19*^*fl/Y*^;rtTA^Tg/+^;TetO-Cre^Tg/0^ (*Pcdh19*^*fl/*Y^;rtTA;Cre) embryos

Dox was administered to pregnant mice (starting at 3.5 dpc) by including in drinking water (2 mg/ml). A sucralose (Splenda powder) at 4 mg/ml final concentration was added to the Dox-containing water, and control (without Dox). A total volume of 250 ml (Sucralose with or without Dox) was transferred to a lightproof bottle and replaced once per week. The embryo stage was determined by the date of the vaginal plug (0.5 dpc).

To confirm the efficiency of Dox-inducible Cre recombinase we performed a diagnostic PCR using the following primers Fwd: CGCTGATTGGCTAGTGCCTG Rev: GAACTTCAGGGTCAGCTTGC. The presence of a ~480 bp band indicated successful *Pcdh19* Exon 1 recombination (Fig. 4B) (Fenno et al. 2014).

### AAV-PHP.eB-EF1α-Cre production and Retro-orbital systemic delivery

The AAV-PHP.eB-EF1a-Cre particles (2.2-3.3 x10^12^ vg/ml) were produced by the Functional Genomics Core Facility (Adelaide University, Australia) using a transgene plasmid pAAV-EF1a-Cre (Addgene plasmid #55636), and PHP.eB capsid (Addgene #103005) and helper (Addgene #112867) plasmids. AAVs were produced at the Functional Genomics South Australia Facility (Adelaide University, Australia) following established Addgene protocols for HEK293T cell transfection, virus harvesting, purification by iodixanol gradient ultracentrifugation, concentration, and viral genome titration.

The AAV-PhP.eB-EF1a-Cre vector were injected intravenously via the retro-orbital route as described (Prabhakar et al., 2021). Briefly, P15 anaesthetized *Pcdh19*-flox mice were injected with 100 ml total volume of AAV at a dose of 1 × 10^11^ vg/mouse or 0.9% saline control via the retro-orbital sinus. One week after the administration, the mice were euthanized and the brains were collected for immunofluorescence and WB analysis (see Fig. 4C) using the methods described below.

### Immunofluorescence assay

The embryonic18.5 dpc mouse brains were fixed in 4% PFA at 4°C overnight. For postnatal brain collection at three weeks, mice underwent cardiac perfusion with 4% PFA, and the brains were dissected and submerged in PFA 4% overnight at 4°C, as previously described (Dawson et al. 2020) (Mincheva-Tasheva et al. 2021). Brains were then cryoprotected in 30% sucrose and frozen in Optimum Cutting 753 Temperature (OCT) embedding medium. Sections (16 µm) were prepared using a Leica CM1900 cryostat. Sections were blocked with 0.1% Triton X-100 and 10% horse serum in PBS for 1 hour at room temperature. Sections were then incubated overnight using with chicken anti-GFP (Abcam, ab13970) 1:400; and rabbit anti-DCX (Abcam, ab18723) 1:500; rabbit anti-Ki67 (Abcam, ab15580) 1:500; mouse anti-NeuN (Sigma, MAB377) 1:400 antibodies at 4°C. Slices were washed three times with PBS1x and incubated with secondary antibody donkey anti-Mouse Alexa 594, anti-Rabbit Cy5 and anti-Chicken 488 (Jackson ImmunoResearch) respectively, for 2 hours at room temperature. Slides were mounted in Prolong™ Gold Antifade with DAPI (Invitrogen #P36931). Images were acquired on a Nikon Eclipse Ti microscope and Nikon Digital Sight DS-Qi1 camera. For each group (experimental vs control), a minimum of three brain sections from at least three experimental and control mice were analyzed. Total DCX fluorescence intensity in the hippocampal dentate gyrus (DG) area was quantified using Fiji (ImageJ) software. Images for each fluorescence intensity comparison were captured using identical microscopy settings. Regions of interest (ROIs) corresponding to the DG area were manually delineated on each section, and the DCX signal was measured as Integrated Density. Background fluorescence was captured from adjacent regions lacking specific staining and subtracted from the measured values. The background-corrected Integrated Density was normalized to the analyzed area, providing a measure of DCX fluorescence intensity per unit area. Data are presented as the average mean intensity per area and expressed as a percentage of control (combined data from *Pcdh19*^*fl/Y*^, WT/Y and *Syn1-Cre* male littermate mice). The total number of Ki67 positive cells in hippocampal DG area per slice were automatically counted using Analyse particles Fiji plugin (n≥3). Data are presented as the average mean intensity per area and expressed as a percentage of control (combined male *Pcdh19*^*fl/Y*^ and *Syn1-Cre* brain samples). Values are reported as mean ± SEM. Statistical analysis was performed using a two-tailed t-test. The *p*-values < 0.05 (*) or < 0.01 (**) were considered significant.

### RT-qPCR analysis

Total RNA was isolated from tissue samples disrupted in Lysing matrix D tubes (MP Biomedicals) containing a Lysis buffer (RNeasy Micro Kit (Qiagen)). After homogenization the supernatant was transferred into the extraction columns from RNeasy Micro Kit (Qiagen), and RNA extraction was performed following the manufacturer’s protocol. cDNA was synthesized using a Super Script III First Strand Synthesis System (Thermo Fisher Scientific). We used predesigned primers from Sigma for *Pcdh19 (*exons 4-5): Fwd: 5′-TTAATCAAAAGCAGCTCCAC, and Rev: 5′-GACATCATTCACAGCAGTATC. As an internal control, a primer set for the housekeeping gene, glyceraldehyde-3-phosphate dehydrogenase (*Gapdh*) was used as follows: Fwd: 5′-AGGTCGGTGTGAACGGATTTG and Rev: 5′-TGTAGACCATGTAGTTGAGGTCA. RT-qPCR was performed using SYBR^®^ Premix Ex Taq™ II (Perfect Real Time, Takara Bio Inc.). SYBR green detection of PCR products was conducted in real time using a MyiQ single-color detection system (Bio-Rad, Hercules, CA, USA). Statistical analysis was performed using an unpaired two-tailed *t*-test. The *p*-values < 0.05 (*) were considered significant.

### Protein extraction and western blotting

Brains were homogenised and extracted in 50mM Tris, 150mM NaCl, 0.2% Triton X-100, 2mM EDTA, 0.01% SDS, 0.1mM Na_3_VO_4_ and 1x Protease inhibitor (Sigma) using Lysing matrix D tubes from MP Biomedicals in a Precellys 3D beater machine. Lysates were separated on Invitrogen Bolt precast 4%–12% polyacrylamide gels, or 7% hand poured SDS-PAGE gels and transferred to PVDF membrane before being blotted. Membranes were blocked in 5% BSA + 5% skim milk in Tris-buffered saline + 0.1% Tween20 (TBST) and rabbit anti-PCDH19 1:650 antibody (ab191198) was incubated with the membrane in 5% BSA in TBST at 4°C overnight. Membranes were developed using Thermo Fisher Scientific Super Signal West Pico PLUS Chemiluminescent Substrate and imaged using a Bio-Rad ChemiDoc. No-Stain Protein Labelling Reagent was used for the total-protein normalisation (Invitrogen, #A44717) prior to primary antibody immunoblot. Statistical analysis was performed using an unpaired two-tailed *t*-test. The *p*-values < 0.05 (*) were considered significant.

### Open Field Test (OFT)

The OFT was conducted as described by (Seibenhener and Wooten 2015). Briefly, 3-week-old mice were acclimatised on Day 1 to the open field arena (44 x 44 cm), for 10 min prior to testing. On Day 2 each mouse was gently placed in the centre of the arena and allowed to freely explore for 10 min. Between trials, the arena was cleaned with 70% ethanol to eliminate olfactory cues. All testing was conducted under consistent lighting and environmental conditions to minimise variability. All parameters presented in Fig. 3C were automatically quantified during the first 5 min of recording using the motion detection and analysis software EthoVision XT developed by Noldus Information Technology. Comparisons between two groups were performed using a nonparametric Mann-Whitney *t*-test.

## Acknowledgments

We thank Prof. Konrad Hochedlinger for providing the C57BL/6-Col1a1tm2(tetO-RNAi:Rps19)Karl Gt(ROSA)26Sortm1(rtTA*M2)Jae/AnuApb mouse line. The authors acknowledge the facilities and the scientific and technical assistance of the South Australian Genome Editing (SAGE) Facility (supported by Phenomics Australia) and the South Australian Health and Medical Research Institute. AAV vectors were generated by Functional Genomics South Australia Core Facility, an NCRIS funded node of Phenomics Australia.

This work was supported by an Australian National Health and Medical Research Council Ideas Grant (APP1129679), the PCDH19 Alliance, the Insieme per la Ricerca PCDH19 – ONLUS, the <u>Firefly Children’s Foundation</u> and Therapeutic Innovation Australia (TIA) Pipeline Accelerator awarded to P. Q. T. and S. M.-T.

## Author contributions

S.M.-T. and P.Q.T.: Conceptualization. S.M.-T., M.S., L.J.R. and S.P.: Investigation. S.M.-T., M.S., J.B. and D.T.P.: Methodology. S.M.-T. and M.S.: Formal analysis. S.M.-T.: Writing-original draft. P.Q.T.: Supervision. S.M.-T. and P.Q.T.: Funding acquisition. All authors: Writing – review and editing.

## Supplementary Materials and Methods

### Droplet digital PCR (ddPCR)

To assay copy number variation (CNV) of the GFP cassette within the *Pcdh19* floxed transgenic mouse, droplet digital PCR (ddPCR) was performed using a Bio-Rad designed assay targeting mGFP for CNV (dCNS372378948). A second assay targeting mouse *Rpp30* (dMmuCNS822293939) was used to normalize samples. DNA (100 ng per sample) was digested with MseI, and run at 4 diluted concentrations (50 ng, 25 ng, 12.5 ng, and 6 ng). Droplet generation was performed on a QX200 (BioRad). The data represented in SuppFig. 1 illustrates the quantification using 25 ng DNA.

## Figure Legends

**Supplementary Figure 1**. Sequences of the donor ssDNAs (3′ and 5′) used to generate *Pcdh19*-cKO-GFP. Digital droplet PCR of GFP copy number in WT, *Pcdh19*^*fl/Y*^ and *Pcdh19*^*fl/WT*^ mice.

**Supplementary Figure 2**. Diagnostic PCR (left panel) amplifying the 5′ LoxP site in Dox-treated (in green) and control-untreated (in black) mice. The right panel illustrates the PCR amplifying only the floxed (reduced in size) *Pcdh19* product in Dox-treated mice. The primer positions are indicated with blue arrows, and red arrowheads indicate the position of the LoxP sites.

## References

Bassani, S., A. W. Cwetsch, L. Gerosa, G. M. Serratto, A. Folci, I. F. Hall, M. Mazzanti, L. Cancedda, and M. Passafaro. 2018. ‘The female epilepsy protein PCDH19 is a new GABAAR-binding partner that regulates GABAergic transmission as well as migration and morphological maturation of hippocampal neurons’, Hum Mol Genet, 27: 1027–38.

Borghi, R., V. Magliocca, S. Petrini, L. A. Conti, S. Moreno, E. Bertini, M. Tartaglia, and C. Compagnucci. 2021. ‘Dissecting the Role of PCDH19 in Clustering Epilepsy by Exploiting Patient-Specific Models of Neurogenesis’, J Clin Med, 10.

Chen, P., F. Chen, Y. Wu, and B. Zhou. 2021. ‘New Insights Into the Role of Aberrant Hippocampal Neurogenesis in Epilepsy’, Front Neurol, 12: 727065.

Dawson, R. E., A. F. Nieto Guil, L. J. Robertson, S. G. Piltz, J. N. Hughes, and P. Q. Thomas. 2020. ‘Functional screening of GATOR1 complex variants reveals a role for mTORC1 deregulation in FCD and focal epilepsy’, Neurobiol Dis, 134: 104640.

de Nys, R., A. Gardner, C. van Eyk, S. Mincheva-Tasheva, P. Thomas, R. Bhattacharjee, L. Jolly, I. Martinez-Garay, I. W. J. Fox, K. S. Kamath, R. Kumar, and J. Gecz. 2024. ‘Proteomic analysis of the developing mammalian brain links PCDH19 to the Wnt/beta-catenin signalling pathway’, Mol Psychiatry.

de Nys, R., R. Kumar, and J. Gecz. 2021. ‘Protocadherin 19 Clustering Epilepsy and Neurosteroids: Opportunities for Intervention’, Int J Mol Sci, 22.

Depienne, C., and E. LeGuern. 2012. ‘PCDH19-related infantile epileptic encephalopathy: an unusual X-linked inheritance disorder’, Hum Mutat, 33: 627–34.

Dibbens, L. M., P. S. Tarpey, K. Hynes, M. A. Bayly, I. E. Scheffer, R. Smith, J. Bomar, E. Sutton, L. Vandeleur, C. Shoubridge, S. Edkins, S. J. Turner, C. Stevens, S. O’Meara, C. Tofts, S. Barthorpe, G. Buck, J. Cole, K. Halliday, D. Jones, R. Lee, M. Madison, T. Mironenko, J. Varian, S. West, S. Widaa, P. Wray, J. Teague, E. Dicks, A. Butler, A. Menzies, A. Jenkinson, R. Shepherd, J. F. Gusella, Z. Afawi, A. Mazarib, M. Y. Neufeld, S. Kivity, D. Lev, T. LermanSagie, A. D. Korczyn, C. P. Derry, G. R. Sutherland, K. Friend, M. Shaw, M. Corbett, H. G. Kim, D. H. Geschwind, P. Thomas, E. Haan, S. Ryan, S. McKee, S. F. Berkovic, P. A. Futreal, M. R. Stratton, J. C. Mulley, and J. Gécz. 2008. ‘X-linked protocadherin 19 mutations cause female-limited epilepsy and cognitive impairment’, Nat Genet, 40: 776–81.

Emond, M. R., S. Biswas, and J. D. Jontes. 2009. ‘Protocadherin-19 is essential for early steps in brain morphogenesis’, Dev Biol, 334: 72–83.

Fenno, L. E., J. Mattis, C. Ramakrishnan, M. Hyun, S. Y. Lee, M. He, J. Tucciarone, A. Selimbeyoglu, A. Berndt, L. Grosenick, K. A. Zalocusky, H. Bernstein, H. Swanson, C. Perry, I. Diester, F. M. Boyce, C. E. Bass, R. Neve, Z. J. Huang, and K. Deisseroth. 2014. ‘Targeting cells with single vectors using multiple-feature Boolean logic’, Nat Methods, 11: 763–72.

Gaitan, Y., and M. Bouchard. 2006. ‘Expression of the delta-protocadherin gene Pcdh19 in the developing mouse embryo’, Gene Expr Patterns, 6: 893–9.

Galindo-Riera, N., S. A. Newbold, M. Sledziowska, C. Llinares-Benadero, J. Griffiths, E. Mire, and I. Martinez-Garay. 2021. ‘Cellular and Behavioral Characterization of’, eNeuro, 8.

Gecz, J., and P. Q. Thomas. 2020. ‘Disentangling the paradox of the PCDH19 clustering epilepsy, a disorder of cellular mosaics’, Curr Opin Genet Dev, 65: 169–75.

Gerosa, L., M. Francolini, S. Bassani, and M. Passafaro. 2019. ‘The Role of Protocadherin 19 (PCDH19) in Neurodevelopment and in the Pathophysiology of Early Infantile Epileptic Encephalopathy-9 (EIEE9)’, Dev Neurobiol, 79: 75–84.

Gerosa, L., S. Mazzoleni, F. Rusconi, A. Longaretti, E. Lewerissa, S. Pelucchi, L. Murru, S. G. Giannelli, V. Broccoli, E. Marcello, N. N. Kasri, E. Battaglioli, M. Passafaro, and S. Bassani. 2022. ‘The epilepsy-associated protein PCDH19 undergoes NMDA receptor-dependent proteolytic cleavage and regulates the expression of immediate-early genes’, Cell Rep, 39: 110857.

Giansante, G., S. Mazzoleni, A. G. Zippo, L. Ponzoni, A. Ghilardi, G. Maiellano, E. Lewerissa, E. van Hugte, N. Nadif Kasri, M. Francolini, M. Sala, L. Murru, S. Bassani, and M. Passafaro. 2023. ‘Neuronal network activity and connectivity are impaired in a conditional knockout mouse model with PCDH19 mosaic expression’, Mol Psychiatry.

Hayashi, S., Y. Inoue, S. Hattori, M. Kaneko, G. Shioi, T. Miyakawa, and M. Takeichi. 2017. ‘Loss of X-linked Protocadherin-19 differentially affects the behavior of heterozygous female and hemizygous male mice’, Sci Rep, 7: 5801.

Herring, C. A., R. K. Simmons, S. Freytag, D. Poppe, J. J. D. Moffet, J. Pflueger, S. Buckberry, D. B. Vargas-Landin, O. Clement, E. G. Echeverria, G. J. Sutton, A. Alvarez-Franco, R. Hou, C. Pflueger, K. McDonald, J. M. Polo, A. R. R. Forrest, A. K. Nowak, I. Voineagu, L. Martelotto, and R. Lister. 2022. ‘Human prefrontal cortex gene regulatory dynamics from gestation to adulthood at single-cell resolution’, Cell, 185: 4428–47 e28.

Homan, C. C., S. Pederson, T. H. To, C. Tan, S. Piltz, M. A. Corbett, E. Wolvetang, P. Q. Thomas, L. A. Jolly, and J. Gecz. 2018. ‘PCDH19 regulation of neural progenitor cell differentiation suggests asynchrony of neurogenesis as a mechanism contributing to PCDH19 Girls Clustering Epilepsy’, Neurobiol Dis, 116: 106–19.

Hoshina, N., E. M. Johnson-Venkatesh, M. Hoshina, and H. Umemori. 2021. ‘Female-specific synaptic dysfunction and cognitive impairment in a mouse model of’, Science, 372.

Kolc, K. L., L. G. Sadleir, I. E. Scheffer, A. Ivancevic, R. Roberts, D. H. Pham, and J. Gecz. 2019. ‘A systematic review and meta-analysis of 271 PCDH19-variant individuals identifies psychiatric comorbidities, and association of seizure onset and disease severity’, Mol Psychiatry, 24: 241–51.

Langford-Smith, A., M. Malinowska, K. J. Langford-Smith, G. Wegrzyn, S. Jones, R. Wynn, J. E. Wraith, F. L. Wilkinson, and B. W. Bigger. 2011. ‘Hyperactive behaviour in the mouse model of mucopolysaccharidosis IIIB in the open field and home cage environments’, Genes Brain Behav, 10: 673–82.

Lim, J., J. Ryu, S. Kang, H. J. Noh, and C. H. Kim. 2019. ‘Autism-like behaviors in male mice with a Pcdh19 deletion’, Mol Brain, 12: 95.

Mathiesen, S. N., J. L. Lock, L. Schoderboeck, W. C. Abraham, and S. M. Hughes. 2020. ‘CNS Transduction Benefits of AAV-PHP.eB over AAV9 Are Dependent on Administration Route and Mouse Strain’, Mol Ther Methods Clin Dev, 19: 447–58.

Mincheva-Tasheva, S., A. F. Nieto Guil, C. C. Homan, J. Gecz, and P. Q. Thomas. 2021. ‘Disrupted Excitatory Synaptic Contacts and Altered Neuronal Network Activity Underpins the Neurological Phenotype in PCDH19-Clustering Epilepsy (PCDH19-CE)’, Mol Neurobiol, 58: 2005–18.

Mincheva-Tasheva, S., C. Pfitzner, R. Kumar, I. Kurtsdotter, M. Scherer, T. Ritchie, J. Muhr, J. Gecz, and P. Q. Thomas. 2024. ‘Mapping combinatorial expression of non-clustered protocadherins in the developing brain identifies novel PCDH19-mediated cell adhesion properties’, Open Biol, 14: 230383.

Mullen, R. J., C. R. Buck, and A. M. Smith. 1992. ‘NeuN, a neuronal specific nuclear protein in vertebrates’, Development, 116: 201–11.

Newbold, S. A., V. Becerra-Espinosa, J. Fabra-Beser, I. W. Fox, C. Llinares-Benadero, E. Stebleva, C. Gil-Sanz, and I. Martinez-Garay. 2026. ‘The intracellular domain of the epilepsy protein PCDH19 regulates spine density in cortical neurons in vivo via Xlr genes’, J Neurosci.

Niu, W., L. Deng, S. P. Mojica-Perez, A. M. Tidball, R. Sudyk, K. Stokes, and J. M. Parent. 2024. ‘Abnormal cell sorting and altered early neurogenesis in a human cortical organoid model of Protocadherin-19 clustering epilepsy’, Front Cell Neurosci, 18: 1339345.

Pederick, D. T., C. C. Homan, E. J. Jaehne, S. G. Piltz, B. P. Haines, B. T. Baune, L. A. Jolly, J. N. Hughes, J. Gecz, and P. Q. Thomas. 2016. ‘Pcdh19 Loss-of-Function Increases Neuronal Migration In Vitro but is Dispensable for Brain Development in Mice’, Sci Rep, 6: 26765.

Pederick, D. T., K. L. Richards, S. G. Piltz, R. Kumar, S. Mincheva-Tasheva, S. A. Mandelstam, R. C. Dale, I. E. Scheffer, J. Gecz, S. Petrou, J. N. Hughes, and P. Q. Thomas. 2018. ‘Abnormal Cell Sorting Underlies the Unique X-Linked Inheritance of PCDH19 Epilepsy’, Neuron, 97: 59–66.e5.

Perl, A. K., S. E. Wert, A. Nagy, C. G. Lobe, and J. A. Whitsett. 2002. ‘Early restriction of peripheral and proximal cell lineages during formation of the lung’, Proc Natl Acad Sci U S A, 99: 10482–7.

Pham, D. H., C. C. Tan, C. C. Homan, K. L. Kolc, M. A. Corbett, D. McAninch, A. H. Fox, P. Q. Thomas, R. Kumar, and J. Gecz. 2017. ‘Protocadherin 19 (PCDH19) interacts with paraspeckle protein NONO to co-regulate gene expression with estrogen receptor alpha (ERα)’, Hum Mol Genet, 26: 2042–52.

Powell, S. B., M. A. Geyer, D. Gallagher, and M. P. Paulus. 2004. ‘The balance between approach and avoidance behaviors in a novel object exploration paradigm in mice’, Behav Brain Res, 152: 341–9.

Rakotomamonjy, J., N. P. Sabetfakhri, S. L. McDermott, and A. Guemez-Gamboa. 2020. ‘Characterization of seizure susceptibility in Pcdh19 mice’, Epilepsia, 61: 2313–20.

Rempe, D., G. Vangeison, J. Hamilton, Y. Li, M. Jepson, and H. J. Federoff. 2006. ‘Synapsin I Cre transgene expression in male mice produces germline recombination in progeny’, Genesis, 44: 44–9.

Robertson, L., D. Pederick, S. Piltz, M. White, A. Nieto, M. Ahladas, F. Adikusuma, and P. Q. Thomas. 2018. ‘Expanding the RNA-Guided Endonuclease Toolkit for Mouse Genome Editing’, CRISPR J, 1: 431–39.

Seibenhener, M. L., and M. C. Wooten. 2015. ‘Use of the Open Field Maze to measure locomotor and anxiety-like behavior in mice’, J Vis Exp: e52434.

Tidball, A. M., W. Niu, Q. Ma, T. N. Takla, J. C. Walker, J. L. Margolis, S. P. Mojica-Perez, R. Sudyk, L. Deng, S. J. Moore, R. Chopra, V. G. Shakkottai, G. G. Murphy, Y. Yuan, L. L. Isom, J. Z. Li, and J. M. Parent. 2023. ‘Deriving early single-rosette brain organoids from human pluripotent stem cells’, Stem Cell Reports, 18: 2498–514.

Weiner, J. A., and J. D. Jontes. 2013. ‘Protocadherins, not prototypical: a complex tale of their interactions, expression, and functions’, Front Mol Neurosci, 6: 4.

Zhu, Y., M. I. Romero, P. Ghosh, Z. Ye, P. Charnay, E. J. Rushing, J. D. Marth, and L. F. Parada. 2001. ‘Ablation of NF1 function in neurons induces abnormal development of cerebral cortex and reactive gliosis in the brain’, Genes Dev, 15: 859–76.

Ziobro, J. M., J. Huang, N. Daddo, E. Margolis, K. Jonatzke, H. Umemori, and J. M. Parent. 2025. ‘Hyperthermic Seizure Susceptibility and Focal Decreases in Parvalbumin-Expressing Cortical Interneurons in a Mouse Model of PCDH19-Clustering Epilepsy’, BioRxiv.

