## Supplementary Figures for "Functional analysis of the epilepsy gene *Pcdh19* using a novel conditional GFP-reporter mouse model"

**Supplementary Figure 1**

| **Oligo** | **Sequence 5’-3’** |
|---|---|
| 3’LoxP ssDNA donor | GGGCTTGCTTTCCTGGAACCCAGTGTTTTGCAGAGGAGAGCTGAAAGGG<br>GATGGTGTACTGGCAGCTGTCTTGTCCTGCTATAACTTCGTATAGCATAC<br>ATTATACGAAGTTATGTAGGGTTATAGTTCACGCGGCCGCGAAGTTCCT<br>ATTCTCTAGAAAGTATAGGAACTTCgccaccgtcgacATGGTGAGCAAGGGCG<br>AGGAGCTGTTCACCGGGGTGGTGCCCATCCTGGTCGAGCTGGACGGCGA<br>CGTAAACGGCCACAAGTTCAGCGTGTCCGGCGAGGGCGAGGGCGATGC<br>CACCTACGGCAAGCTGACCCTGAAGTTCATCTGCACCACCGGCAAGCTG<br>CCCGTGCCCTGGCCCACCCTCGTGACCACCCTGACCTACGGCGTGCAGT<br>GCTTCAGCCGCTACCCCGACCACATGAAGCAGCACGACTTCTTCAAGTC<br>CGCCATGCCCGAAGGCTACGTCCAGGAGCGCACCATCTTCTTCAAGGAC<br>GACGGCAACTACAAGACCCGCGCCGAGGTGAAGTTCGAGGGCGACACC<br>CTGGTGAACCGCATCGAGCTGAAGGGCATCGACTTCAAGGAGGACGGC<br>AACATCCTGGGGCACAAGCTGGAGTACAACTACAACAGCCACAACGTCT<br>ATATCATGGCCGACAAGCAGAAGAACGGCATCAAGGTGAACTTCAAGA<br>TCCGCCACAACATCGAGGACGGCAGCGTGCAGCTCGCCGACCACTACCA<br>GCAGAACACCCCCATCGGCGACGGCCCCGTGCTGCTGCCCGACAACCAC<br>TACCTGAGCACCCAGTCCGCCCTGAGCAAAGACCCCAACGAGAAGCGC<br>GATCACATGGTCCTGCTGGAGTTCGTGACCGCCGCCGGGATCACTCTCG<br>GCATGGACGAGCTGTACAAGTAACGACTGTGCCTTCTAGTTGCCAGCCA<br>TCTGTTGTTTGCCCCTCCCCCGTGCCTTCCTTGACCCTGGAAGGTGCCAC<br>TCCCACTGTCCTTTCCTAATAAAATGAGGAAATTGCATCGCATTGTCTGA<br>GTAGGTGTCATTCTATTCTGGGGGGTGGGGTGGGGCAGGACAGCAAGG<br>GGGAGGATTGGGAAGACAATAGCAGGCATGCTGGGGATGCGGTGGGCT<br>CTATGGGAAGTTCCTATTCTCTAGAAAGTATAGGAACTTCGCGGCCGCA<br>GAGAGAGAGCTCTGAGAGTTTATGATAAGAAACGG |
| 5’LoxP ssDNA donor | G*C*T*A*CCGCTCACCATGCCCGGCTTGCCCAGAGGGCACTGGCATTCA<br>CTTCCCTAGCCCCTATAACTTCGTATAGCATACATTATACGAAGTTATCG<br>CGGGCTGCGGCGCAGTGTCCTCCCTGGCTTTCCAGCTGAGCCGATTCATC<br>TGAG*A*C*C*C<br><br>*Nucleotides with phosphorothioate modification. |

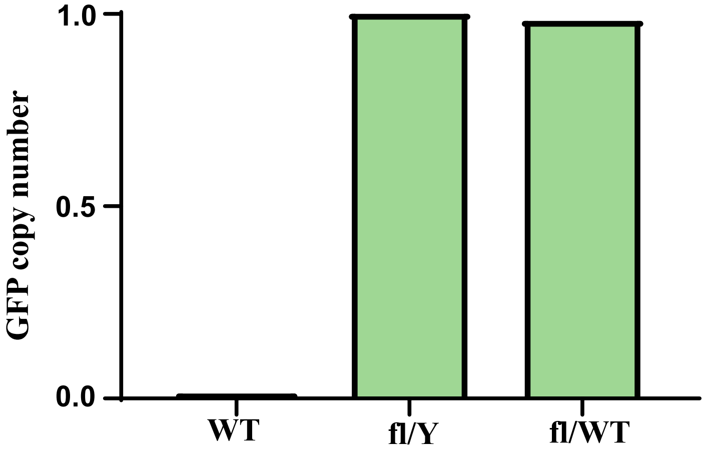


**Supp.Fig.1** Sequences of the donor ssDNAs (3’ and 5’) used to generate *Pcdh19*-cKO-GFP. The graph illustrates the digital droplet PCR of GFP copy number in WT, *Pcdh19*fl/Y and *Pcdh19* fl/WT mice (n=1).

**Supplementary Figure 2**

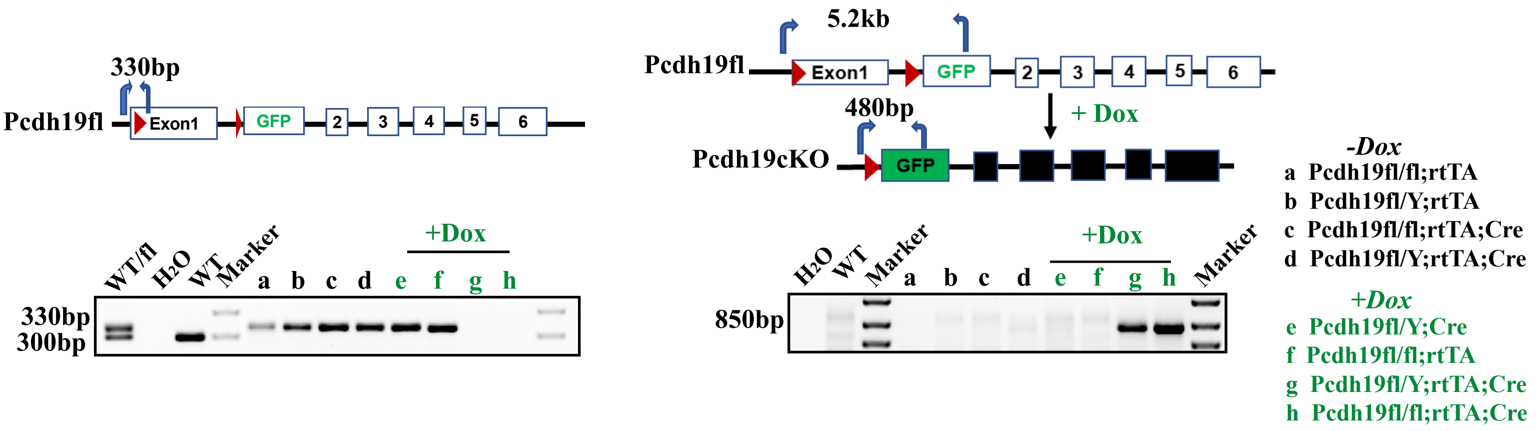


**Supplementary Figure 2**

Diagnostic PCR (left panel) amplifying the 5'LoxP site in Dox-treated (in green) and control -untreated (in black) mice. The right panel illustrates the PCR amplifying only the floxed (reduced in size) Pcdh19 product in Dox-treated mice. The primer positions are indicated with blue arrows and red arrowheads indicate the position of the LoxP sites.
